# Niche specificity, polygeny, and pleiotropy in herbivorous insects

**DOI:** 10.1101/2021.03.12.435098

**Authors:** Nate B Hardy, Matt Forister

## Abstract

What causes host-use specificity in herbivorous insects? Population genetic models predict specialization when habitat preference can evolve and there is antagonistic pleiotropy at a performance-affecting locus. But empirically for herbivorous insects, host-use performance is governed by many genetic loci, and antagonistic pleiotropy seems to be rare. Here, we use individual-based quantitative genetic simulation models to investigate the role of pleiotropy in the evolution of sympatric host-use specialization when performance and preference are quantitative traits. We look first at pleiotropies affecting only host-use performance. We find that when the host environment changes slowly the evolution of host-use specialization requires levels of antagonistic pleiotropy much higher than what has been observed in nature. On the other hand, with rapid environmental change or pronounced asymmetries in productivity across host species, the evolution of host-use specialization readily occurs without pleiotropy. When pleiotropies affect preference as well as performance, even with slow environmental change and host species of equal productivity, we observe fluctuations in host-use breadth, with mean specificity increasing with the pervasiveness of antagonistic pleiotropy. So, our simulations show that pleiotropy is not necessary for specialization, although it can be sufficient, provided it is extensive or multifarious.

## Introduction

Most species of herbivorous insects use as hosts only a small fraction of the plant species they encounter in their environment (Forister et al., 2015; Futuyma & Moreno, 1988), despite the obvious advantages of using many hosts and having a broader resource base (Hardy et al., 2020; Rainey et al., 2000; Smith & Hoekstra, 1980). For the most part, theorists have explained the evolution of this specificity as a consequence of antagonistic pleiotropy (Egas et al., 2004; Rueffler et al., 2004): an allele that confers high performance on one host confers low performance on another. Mathematical models show that if host-use performance is determined by one gene with antagonistic pleiotropy, and if populations can evolve host-use preferences, then the evolution of host specialization is all but inevitable, at least for panmictic populations in static environments (Ravigné et al., 2009).

But in two key aspects, this theory bears little resemblance to what we know about herbivorous insects. First, in herbivorous insects, most host-specific performance phenotypes, such as the ability to develop on a diet of any particular plant species, seem to be controlled by many genes, not by one or even a few (Egan et al., 2015; Vertacnik and Linnen 2017). Second, empirical studies of polygenic local adaptation (reviewed by Savolainen et al. 2013) show that antagonistic pleiotropy tends to be rare in comparison to conditional neutrality; alleles tend to affect performance in one environment but not another. As a striking example, in the Melissa blue butterfly, *Lycaeides melissa*, more than 99% of alleles with host-dependent performance effects are conditionally neutral (Gompert et al. 2015). Although linkage disequilibrium can also cause negative genetic covariances (Via & Lande, 1985), conditionally neutral alleles might seem less likely than pleiotropic alleles to drive the evolution of host-use specialization. Moreover, it is not obvious what influence a few – or many – pleiotropic loci might have in the evolution of a multi-dimensional quantitative trait such as host-use. Here, we use forward-time individual-based quantitative genetic simulations to address this simple question: do simple genetic trade-offs matter in a polygenic world? More specifically, how might pleiotropy affect the evolution of specialization, when preferences can evolve along with performance?

Our models build on much previous work. The theory of the evolution of niche specialization could fill volumes. Ravigné et al. (2009) classify more than seventy studies by five factors: (1) the specific research question addressed, (2) the type of population regulation, (3) the shape of the localadaptation trade-off, (4) whether or not habitat choice was allowed to evolve, and (5) if so, what the mechanism was for the evolution of choice. Our question is how the genetic architectures of habitat preference and performance affect the evolution of niche specialization. In that respect, our work departs from what has come before, almost all of which has focused on the emergence or maintenance of local-adaptation polymorphisms with single-locus architectures for performance and/or preference. Of the few previous studies that take a quantitative genetic approach (e.g. Via and Lande 1985; van Tienderen 1991, 1997), none examine the joint evolution of habitat preference and performance. Also, because our models are individual-based, population regulation and adaptive trade-offs are not as they are in population-based models. Concretely, in our models, population size is something that emerges via the balance of individual mortality, viability, and fecundity. Likewise, adaptive trade-offs – that is, quantitative genetic covariances – are not fixed, but rather emerge via genetic linkage and the pleiotropic effects of specific mutations, and evolve via subsequent mutation, recombination, genetic drift, and natural selection. As for the mechanisms of habitat choice, our models resemble so-called “two-allele” models (Felsenstein 1981), although in this case many more than two alleles are involved. Habitat choice evolves at least somewhat independently from local adaptation, and preference alleles affect proclivity for a specific environment, rather than attachment to wherever they happen to have come from, or whatever would be ideal for their performance.

Although we are primarily interested in the evolution of niche breadth, since our models allow for ecologically-driven population genetic divergence without geographic isolation, they also speak to sympatric speciation, the theory of which is based on many of the same assumptions as that of niche specialization, and was reviewed by Kirkpatrick and Ravigne (2002) and Bolnick and Fitzpatrick (2007). Previous sympatric speciation models vary by (1) the causes of disruptive selection, (2) how individuals select mates, (3) whether selection acts directly or indirectly on mating traits, and (4) the genetic basis of mating traits. In our model, disruptive selection stems from resource heterogeneity, mating is assortative, and selection acts indirectly on the mating trait, namely host-preference (mating takes place on the host), as fitness is only affected by a host-dependent performance phenotype. To our knowledge, our model is the first to have this particular combination of features, or to allow for conditional neutrality at quantitative trait loci.

Our models show that for pleiotropy to drive quantitative specialization, it needs to be quite extensive. Therefore, we also consider what else might drive specialization in sympatry. For direction, we turn to the foundational quantitative genetics work of Via and Lande (1985). With an analytical approach, they identified three conditions for specialization, to wit, there is (1) no gene flow between habitats, (2) insufficient genetic diversity for the relevant niche traits, or (3) a genetic correlation across habitats equal to ± 1, which would be true if phenotypes are monogenic. Otherwise, populations evolve high-performance generalism. Nevertheless, the *rate* of adaptation can vary across habitats. For example, Via and Lande (1985) show that if habitat types vary in frequency, adaptation will be faster and more direct in the more common habitat, something which has also been demonstrated with phenotype-free population genetic models (Whitlock 1996; Kawecki et al. 1997). Moreover, until the population evolves a near-optimal phenotype in the common habitat, the evolutionary response in the rare habitat is largely correlational. And if genetic covariances are negative, this can temporarily increase maladaptation to the rare habitat. To sum up: barring monogeny, monophorphism, and allopatry, given enough time, habitat generalism will evolve, but along the way negative genetic covariances and asymmetries in productivity between habitats increase the odds of temporary specificity.

These insights are contingent on several model assumptions: that (1) habitat-specific phenotypic optima are fixed, (2) habitat preference does not evolve, (3) selection is weak and (4) population size is infinite, so that (5) the matrix of genetic variances and covariances is fixed. Our models relax each of assumptions. Therefore, they provide a more general – although less elegant – view of quantitative niche breadth evolution.

## Methods

Simulations were performed with SliM 3 (Haller & Messer, 2019). In this framework, models are specified in the Eidos programming language. Eidos codes are available from Zenodo (https://doi.org/10.5281/zenodo.6819401). For a list of model parameters and variables see table 1.

**Table 1.** Parameter definitions and values; variable definitions and intervals.

Parameters
| Symbol | Definition | Values |
| --- | --- | --- |
| $\mu$ | Mutation rate | {1e-7; 1e-9} |
| $\chi$ | Recombination rate | {1e-4; 1e-6} |
| $\lambda$ | Mean and variance of Poisson-distributed fecundity | {1.5} |
| $n$ | Number of host species | {2} |
| $\sigma^2$ | Allele effect variance | {0.6} |
| $\kappa$ | Pleiotropic effect covariances | {-0.9$\sigma^2$, 0, 0.9$\sigma^2$} |
| $M$ | Mix of pleiotropic to conditionally neutral mutations | {0, 2<sup>-3</sup>, 2<sup>0</sup>, 2<sup>3</sup>, 2<sup>6</sup>} |
| $\rho$ | Mix of preference to performance mutations | {0, 0.1, 1} |
| $\omega$ | Gaussian fitness function variance | {3.0} |
| $\delta_o$ | Per-generation rate of change in performance optima | {0.005, 0.05, 0.1} |
| $K_j$ | Carrying capacity of host species $j$ | {500} |
| $pK_1$ | Proportional carrying capacity of host 1 | {0.05, 0.25, 0.5} |

Variables
| Symbol | Definition | Intervals |
| --- | --- | --- |
| $h_j$ | Host species $j$ | {$h_1, h_2$} |
| $N$ | Total population size | $0 \leq N$, with a soft maximum of $\Sigma K_j$ |
| $N_j$ | Population size on host species $j$ | $0 \leq N_j$, with a soft maximum of $K_j$ |
| $z_{ij}$ | Performance genotype value of individual $i$ on $h_j$ | $z_{ij} \in \mathbb{R}$ |
| $y_1$ | Genotype value for preference for $h_1$ | $y_1 \in \mathbb{R}$ |
| $O_j$ | Performance optimum for $h_j$ | $0 \leq O_j \leq \delta_o$ * post-burnin generations |
| $H$ | Population heterozygosity | $0 \leq H \leq 1$ |
| $H'$ | Shannon's diversity of host-use | $0 \leq H' \leq \ln(n)$ |
| $v(z_{ij})$ | Genotype-environment match viability | $0 \leq v(z_{ij}) \leq 1$ |
| $W$ | Mean population fitness across host species | $0 \leq W \leq 1$ |
| $F_{ST}$ | Index of neutral genetic subpopulation divergence | $0 \leq F_{ST} \leq 1$ |
| $\delta P$ | Difference in mean $y_i$ between subpopulations | $0 \leq \delta P$ |
| $\delta W$ | Specialization index | $0 \leq \delta_w \leq 1$ |

### *Initialization* (fig1a)

At the start of each simulation, we initialize a population composed of *n*=2 subpopulations, each of which is composed of *N_j_* hermaphroditic individuals. We imagine that these individuals are insect herbivores, and that each subpopulation is associated with host species *h_j_*. We give each individual the same diploid 100 kb, one-chromosome genome. This initial genome is an empty container for future mutations. It is divided into ten 5-kb coding regions interspersed with ten 5-kb non-coding regions. Host-preference genes are mixed in with host-performance genes in the coding regions of the genome, and in some models the same loci affect both traits. Of course, real herbivorous insect genomes are much larger and have multiple chromosomes. Working with smaller genomes is easier computationally, but increases linkage disequilibrium. To correct for that, we ran some simulations with a recombination rate, *χ*, of 1e-4 events per base pair (bp) per generation. At that rate, we expect about ten crossing-over events per genome per generation, which corresponds roughly to as much genetic shuffling as would be typical for a real herbivorous insect species (Wilfert et al., 2007). This assumes that preference and performance alleles are randomly distributed across the genome. To assess the impact of tighter linkage, we also ran simulations with *χ*=1e-6.

In most simulations, the mutation rate, *μ*, is 1e-7 per bp per generation, yielding an expected 0.01 non-neutral mutations per individual per generation. This is an order of magnitude more than what we would expect based on empirical measures of mutation rates (~6e-9 per bp per generation; Liu et al. 2016) and average gene size (1,346 bp; Xu et al. 2006), and if we assume that quantitative host-use performance and preferences are each controlled by ~100 genes (Vertacnik and Linnen 2017). But an elevated per-individual mutation rate compensates for the fact that our simulated populations are smaller than most real ones. Four types of mutations can occur: (1) neutral mutations in non-coding regions, (2) pleiotropic mutations, that is, mutations at quantitative trait loci (QTLs) that affect performance on each host species, and in some models also affect host preference, (3) conditionally neutral mutations at QTLs that affect performance only on host *h_j_*, and (4) conditionally neutral mutations at QTLs that affect preference for host *h_j_*.

After initialization, we simulate many generations following the life cycle shown in fig. 1.

**Figure 1.**
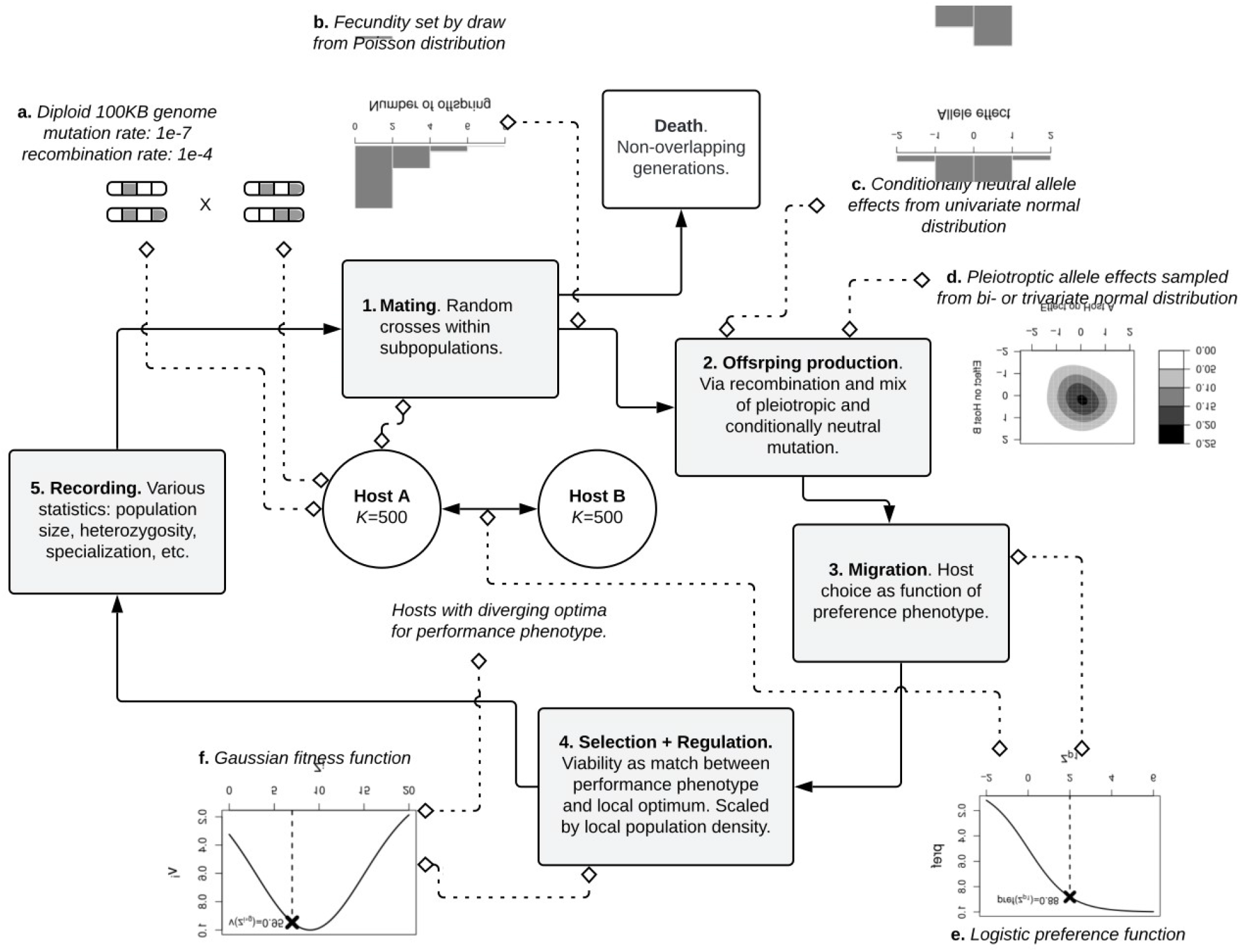
Simulated herbivore life cycle. Each generation consists of (1) a round of random mating followed by (2) offspring production, in which parent genomes crossed and mutated, and then offspring are added to the parental host subpopulation. After reproducing, parents die. The environment consists of two host plant species. Offspring chose to stay on their natal host species, or (3) migrate to the other host species. After a host is chosen, (4) viability is first determined by the match between an individual’s performance phenotype and their host environment, and then scaled by local subpopulation density.

#### 1. Mating

At the start of each generation, each surviving individual randomly selects as a mate one individual from their subpopulation. This reflects the tendency of herbivorous insects to mate on or near their host plants. The fecundity of the choosing individual is determined by a draw from a Poisson distribution with a mean and variance (*λ*) of 1.5 (fig. 1b). The drawn number of crosses between a mating pair (chooser and chosen) are then added to the subpopulation. Note that choosers can also be chosen as mates by other individuals in their subpopulation; in theory, an individual could be randomly chosen as the mate of every other individual in their subpopulation, but the chances of being involved in more than two or three matings is small. As is typical for herbivorous insects, generations are discrete and non-overlapping; after mating and producing offspring, parents die.

#### 2. Offspring production

This entails mutation and recombination of parental genomes. When a mutation occurs at a conditionally neutral performance or preference QTL, an allele effect is drawn from a univariate normal distribution with a mean of zero and a variance (*σ^2^*) of 0.6 (fig. 1c). When a mutation occurs at a pleiotropic performance QTL, its effects are drawn from an bivariate normal distribution (fig. 1d) with means of zero, variances of 0.6, and symmetrical covariances, *κ*, which varied over simulations {-0.9*σ^2^*, 0, 0.9*σ^2^*}. With these distributions of allele effects, on average, simulated populations adapt to changing host environments via promotion of on the order of a few tens of alleles. As previously mentioned, we also examined models in which pleiotropic mutations affected host preference as well as performance in each host environment. For these simulations pleiotropic allele effects are drawn from a tri-variate normal distribution with means of zero, variances of 0.6, the same range of performance trait covariances as for the two-dimensional models, and zero covariances between the preference and performance traits. Our main goal was to see how host-use evolution depends on the ratio of pleiotropic to conditionally-neutral alleles governing host-use performance, and (in some cases) also preference. Therefore, across simulations, we varied this mutational mix, *M*, {0, 2^−3^, 2^0^, 2^3^, 2^6^}.

The relative rates of preference and performance mutations, *ρ*, could affect the odds of population divergence and specialization. Indeed, if mutations that affect host preferences are common, they could be an adaptive hazard, that is, a genetic architectural liability that increases the risk of maladaptation (Lynch 2011). For models with performance-only pleiotropies, we considered two values for *ρ* spanning a range of plausibility {0.1, 1}; this was based on our reading of the relevant gene expression research literature (e.g., Andersson et al. 2014; Wu et al. 2016; Yu et al. 2016; Tan et al. 2019). To get a sense for the sensitivity of some model results to the effects of performance-affecting alleles, we also ran some simulations with *ρ*=0. For models with performance-and-preference pleiotropies, we considered only a 1:1 ratio of conditionally-neutral mutations affecting preference and performance.

#### 3. Migration and habitat choice

Each newly-produced individual selects a host species. At the start of a simulation, all individuals have a preference genotype value, *y_1_*, of zero, and the probability of choosing *h_1_* is 0.5. Then, as the simulation progresses, host preference evolves as a function of the sum of allele effects at preference-determining loci (fig. 1e). Concretely, the probability that an individual will choose *h_1_* over *h_2_* is given by a logistic function, *P*(*y*_1_) = - 1 / (1 + exp(−1 * *y*_1_). This allows from some geneflow as mean preference phenotypes slowly approach either of the boundary states, 0 ≤ *P*(y_1_) ≤ 1.

#### 4. Selection and regulation

The next step in the life cycle is hard selection. An individual’s viability, that is, their probability of surviving long enough to mate, is determined by two factors: (1) the match between their performance genotype and host-environment (fig. 1f), (2) local subpopulation density. An individual’s performance genotype for host *h_j_* is the sum of allele effects across pleiotropic loci and the conditionally neutral loci affecting performance only that host. We assume no epistasis or phenotypic plasticity. Performance phenotypes map to viability with a standard Gaussian fitness function:

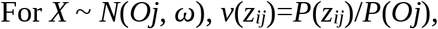

where *z_ij_* is an individual’s genotype value for performance on *h_j_*, *v*(*z_ij_*) is the viability of an individual with *z_ij_* on *h_j_*, *O_j_* is the optimal value for the performance phenotype on *h_j_*, *ω* is the variance {3.0}, that is, the steepness of the selection gradient (smaller variances map to steeper gradients), and division by *P*(*O_j_*) sets the viability of a perfect match to one. The force of selection on a population depends on both *ω* and the rate, *δ_O_*, at which host optima move away from their initial values {0.005, 0.05, 0.1}.

After genotype-environment matching, individual viability is re-scaled by local subpopulation density. Specifically, each host *h_j_* has a nominal carrying capacity, *K_j_*, which is set to 500 individuals. Viabilities are re-scaled by the ratio *K_j_* to the current subpopulation size *N_j_*. By capping this density scaling factor at 1.0, we make selection hard; when *N_j_* > *K_j_*, the viability of all individuals on a host species is reduced, but low density (*N_j_* < *K_j_*) is not enough to drive demographic growth of maladapted genotypes. So, *N* has a soft upper limit equal to the sum of the nominal carrying capacities of host species. If selection is strong enough that the product of mean viability and fecundity is < 1, the population will shrink, and if that product remains < 1, the population will go extinct.

After selection, the life cycle again starts again.

#### The arc of a simulation

At the start of each simulation the optimal value for each performance phenotype *O_j_*, is set to zero, which matches the starting genotype value of each individual in the population. Thus, herbivore populations start off well-adapted to their host environments. *O_j_* stays at zero for a 1,000 generation burn-in period in which the simulated population accumulates genetic diversity. Then, in each of 1,000 subsequent generations, *O_j_* shifts by a random draw from an exponential distribution with mean *δ_O_*. Such gradual and directional changes approximate a variety of real-world situations, for example the effects of global warming, land-use intensification, or biological invasion, and allow host-use adaptation to proceed via incremental changes to mean genotype values.

#### Model variants and statistics

We examined two main models. In Model 1, pleiotropic mutations affect only performance. In Model 2, pleiotropic mutations affect both host-use performance and preference. For both models, we explore various combinations of values for *δ_O_, M, p, χ*, and *κ* (Table 1). For Model 1, we also examined the effects of asymmetries in the productivity of host environments; fixing the global carrying capacity to 1,000, and letting the proportional carrying capacity of *h_1_, pK_1_* vary {0.05, 0.25, 0.5}.

For different perspectives on the genetic states of the population at the end of each simulation under Model 1, we recorded the mean values over a simulation for these statistics: (1) *N*, total population size across hosts, (2) *H*, heterozygosity, (3) *H’*, host-use diversity of the herbivore population, characterized with Shannon’s diversity index, (4) *W*: mean population fitness across host-associated subpopulations; (5) (5) *F_ST_*: the standard ratio of between-subpopulation neutral genetic variance to total neutral genetic variance; (6) *δP*, the difference in mean preference phenotypes between host species, (7) *δW*, a specialization index that measures subpopulation divergence, defined as the mean difference between an individual’s fitness on their host species and what their would have been on other host species, scaled by the overall mean fitness of the population (ranging from zero to one, with smaller values denoting more generalist populations). Below, we focus on whichever statistics we found most revealing.

Before we look at any results, here is some guidance on how to interpret the pleiotropic effect covariances. After the burn-in period, the optimal value for the host-performance phenotype for each host species, *O_i_*, increases by *δ_O_* each generation, and the mean performance trait values in the herbivore population tend to lag behind these diverging optima. Thus, pleiotropic performance mutations drawn from a bi-variate normal distribution with positive covariances tend to move the phenotype either towards or away from the diverging host-use optima (fig. S2), in other words, these are “general vigor” or “general weakness” alleles (Fry 1996). In the former case, pleiotropy can help generalists solve the problem of adapting to a heterogeneous host environment. In contrast, with negative pleiotropic covariances, alleles that increase performance on one host tend to decreases it on others, that is, antagonism abounds. With covariances of zero, positive and antagonistic pleiotropy are equally likely. For Model 2, the covariances between preference and performance trait effects are zero; in other words, alleles that affect performance also affect preference, but positive and negative relations are equally likely between effects on preference and the effects on each performance phenotype.

## Results and Discussion

### Model 1. Performance pleiotropies

Let us first consider what happens when the changes in host-performance optima are slow (*δ_O_*=0.005). Figs. 2a-d show an example run, where there is an even mix of pleiotropic and conditionally neutral mutation (*M*=1), pleiotropic covariances are zero (*κ*=0), the recombination rate is what is expected with a uniform distribution of QTLs across the genome (*χ*=1e-4), and performance and preference mutations occur at an equal rate (*ρ*=1). Population genetic summary statistics vary from one generations to the next, but without strong directional trends; *δW* stays near zero (no specialization), *W* stays near unity (high fitness), *N* hovers around 2*K_j_*, and *H’* stays near its maximum value. The population maintains high-performance generalism. Figs. 3a-d summarize batches of 20 simulations for each combination values for *M*, and *κ* given in Table 1, and with *ρ*=1.0. Each plot shows how a population variable tends to change with increasing amounts of pleiotropy in the mutational mix (*M*), as well as how those changes depend on the covariances of pleiotropic effects (*κ*). Unless antagonistic pleiotropy is many times more common than conditional neutrality (*M*>>1), populations maintain high-performance generalism. *W* is near unity, *H’* is near its maximum value, and *δW* is near zero. Subpopulations do divergence some in host preference, with *δP* ~ 0.2, but this is firmly on the non-choosy end of the spectrum. Even with pervasive and antagonistic pleiotropy (*M*=2^6^; *κ*=-0.9*σ^2^*) the evolution of host-use specialists is unlikely. Moreover, high levels of antagonistic pleiotropy tend to reduce mean fitness; thus insofar as mutational covariances are evolvable, selection will be against antagonism. Antagonistic pleiotropy is even less likely to drive specialization or strong host preferences when preference mutations are less common than performance mutations, *ρ*=0.1, (figs. S2a-d). Lowering the recombination rate between performance alleles (*χ*=1e-6) increases the scope for specialization via pleiotropy, but even so, only when *ρ*=1.0, M>1 and *κ*=-0.9*σ^2^* (figs. S3a-d). In summary, with slow changes in performance optima, barring exceptionally tight linkage, negative genetic covariances, and high preference mutation pressure, quantitative specialization via any amount of pleiotropy is unlikely.

**Figure 2.**
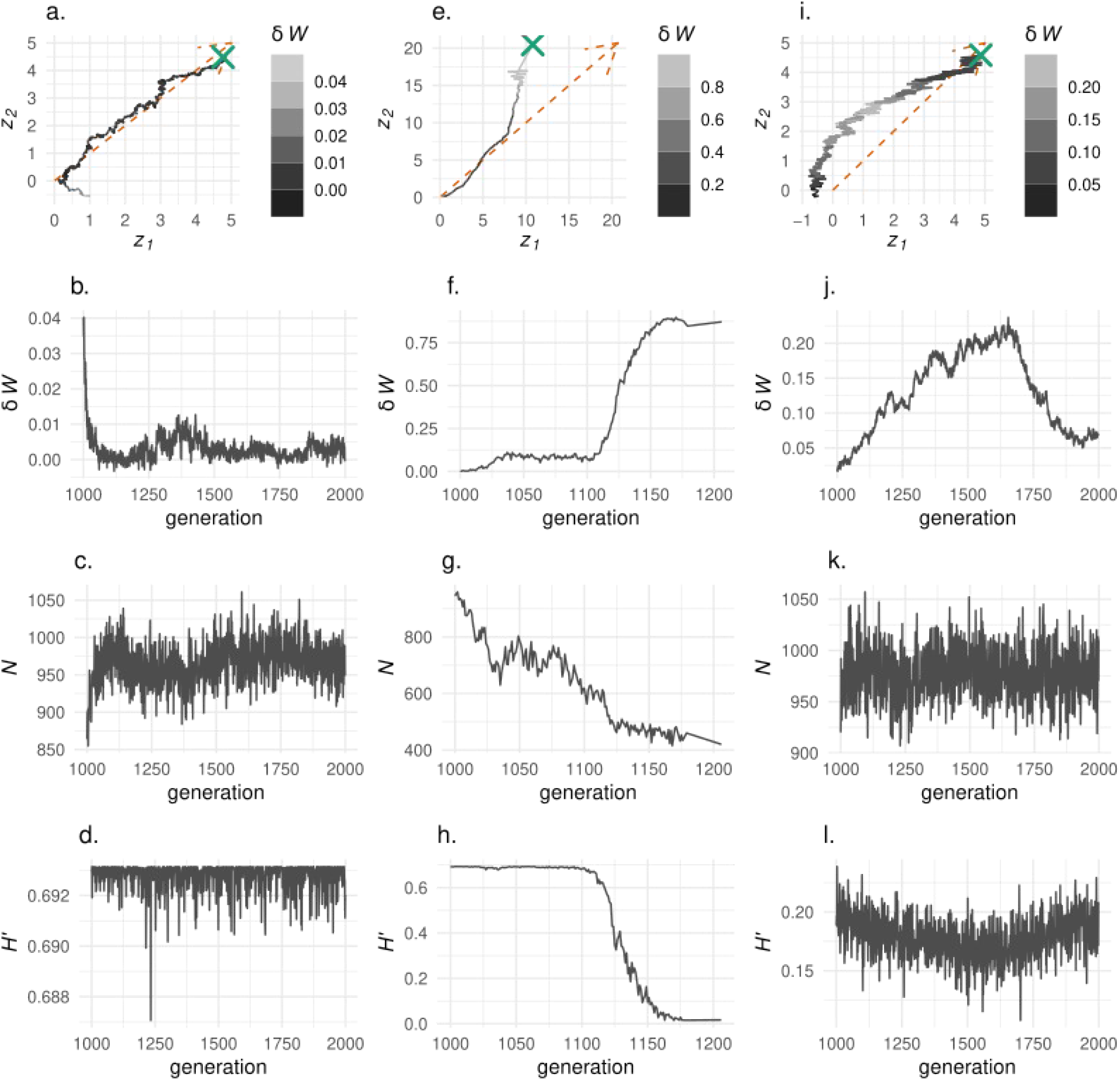
Example runs under Model 1. The left column (a–d) shows a simulation in which host species are equally productive (*K_1_*=*K_2_*), preference mutations are as common as performance mutations (*ρ*=1), and in which optimal performance values change slowly (*δ_O_*=0.005). The middle column (e–h) shows a simulation in which *K_1_=K_2_, ρ*=1, and optimal performance value change rapidly (*δ_O_*=0.1). The right column (i–l) shows a simulation in which *K_1_<K_2_* and optimal performance values change slowly. The top row shows changes in population mean values for performance traits (*z_1_* and *z_2_*), over 1000 generations after the burn-in period. The color of the tracing line changes with a population’s mean host-performance specificity; lighter shades of gray denote higher specificity. The dashed orange arrow shows the concomitant change in host performance optima (*O_j_*). The green ‘X’ denotes the population mean at the end of the simulation.

**Figure 3.**
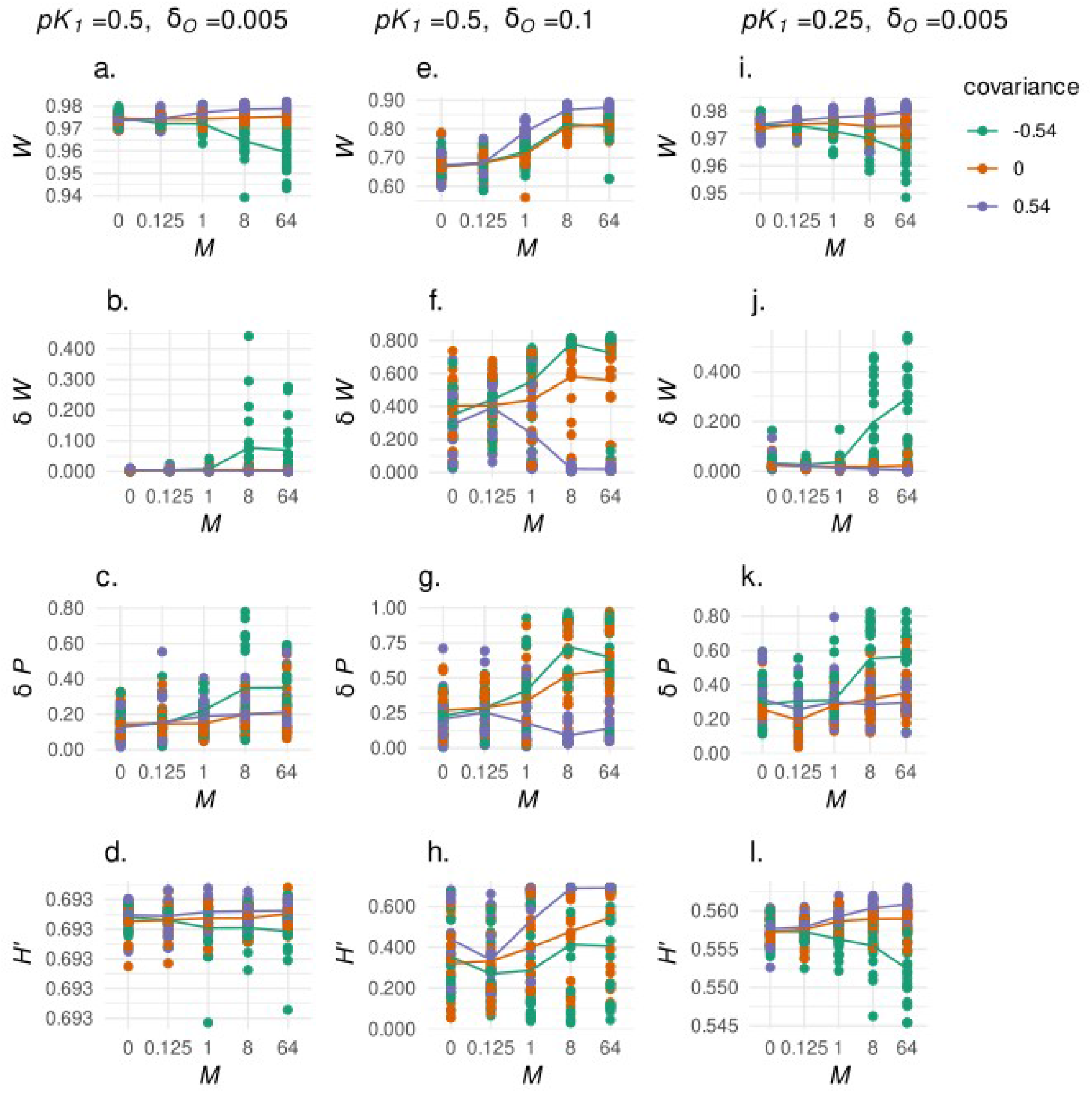
Summary of runs under Model 1 with *ρ*=1. *M* is the ratio of pleiotropic to conditionally neutral mutation. Green lines are for pleiotropic covariances of −0.54, orange lines are for covariances of zero, and purple lines are for covariances of 0.54. All plots show mean population states over 1000 generations after the burn-in period. The left column (a–d) shows population states when host species are equally productive (*K_1_*=*K_2_*) preference mutations are as common as performance mutations (*ρ*=1), and in which optimal performance values change slowly (*δ_O_*=0.005). The middle column (e–h) is for simulations in which *K_1_=K_2_, p*=1, and optimal performance value change rapidly (*δ_O_*=0.1). The right column (i–l) is for simulations in which *K_1_<K_2_ (pK_1_* = 0.25) and optimal performance values change slowly.

Now consider what happens when performance optima diverge rapidly. Figs. 2e-h show an example run in which *M*=0 (there is no pleiotropy), *δ_O_*=0.1, and *ρ*=1. See the population veer away from the joint optimal performance vector; see the rise of *δW* and the fall of *N* and *H’*. We stopped the simulation at generation 1206, at which point the *N*_1_ = 0, and the population is composed exclusively of *h_2_* specialists. Figs. 3e-h, summarize a batch of runs in which we vary *M* and *κ* while *δ_O_* = 0.1, *χ*=1e-4, and *ρ*=1.0. Even when pleiotropy is rare and neutral, populations tend to evolve some level of host-use specialization; *δW* ~ 0.5, and *H’* is about half of its maximum value. The effect of *M* depends on *κ*; increasing *M* increases *δW* and *δP* when *κ=-0.9σ^2^* and decreases it when *κ* = 0 or 0.9*σ^2^*. Also note that regardless of the value of *κ*, increasing the amount of pleiotropy in the mutational mix (*M*) increases mean viability (*W*); in one way or another, depending on its sign, pleiotropy helps the metapopulation adaptively keep pace with the changing host environment. On the one hand, positive pleiotropies provide a double-shot of the adaptive genetic diversity that is depleted by strong and sustained directional selection. On the other hand, antagonistic pleiotropies can push for the evolution of strong host preference, which increase rates of host-use adaptation by effectively homogenizing the host environment to which a genotype will likely be exposed.

Decreasing *χ* further increases the odds of specialization with little pleiotropy (figs. S3e-h), whereas decreasing *ρ* does the opposite (fig. S2e-h). In summary, when the host environment changes rapidly, the odds of host-use specialization increase across genetic architectures. But antagonistic pleiotropy is not necessary. Specialization can also evolve via tight linkage of conditionally neutral alleles, or via high preference mutation pressure.

Another thing that increases the odds of specialization is asymmetry in the productivity of host environments. Figs. 2i-l show an example run for the case in which *pK_1_* = 0.05 – that is *K_1_* accounts for 5% of the total carrying capacity across hosts – and where *M*=0, *ρ*=1, *δ_O_*=0.005, and *χ*=1e-4. As per the classical theory of Via and Lande (1985), performance evolution is initially dominated by adaptation to the more common host, *h_2_*. But with the evolution of divergent specialization between subpopulations, the population’s evolutionary path bends towards the joint optimum vector. In this particular simulation, the population passes through this phase of specialization and begins to coalesce and regeneralize. In other runs (not shown), subpopulations stay specialized; the population’s evolutionary path runs parallel to and above (with *z_2_* on the y-axis) the joint optimal performance vector for the duration of the 1,000 generation observation period. But the fact that populations do occasionally coalesce resonates with previous work demonstrating the importance of gene flow from source subpopulations for adaptation in marginal sink subpopulations (Holt et al. 2003).

Figs. 3i-l summarize a batch of runs with *pK_1_* = 0.25, *ρ* =1, *χ* = 1e-4, and *δ_O_* =0.005, varying *M* and *κ*. Here again we see that even without pleiotropy populations can evolve intermediate levels of host-use specificity (*δW* ~ 0.1), although to a lesser extent then when *δ_O_*=0.1 or *χ*=1e-6. And here again, it is not until *M* > 1 that antagonistic pleiotropy really begins to affect specificity, in this case causing a precipitous decline in *H’*. When host species vary in productivity, pervasive antagonistic pleiotropy tends to push for specialization on the more productive host.

### Model 2. Preference-and-performance pleiotropies

Fig 4 shows an example run with *χ* = 1e-4, *ρ* = 1, *κ* = - 0.9*σ*^2^/2, *M* = 2^−3^, and *δ_O_* = 0.005. Note that there are stretches in which both host environments are occupied by non-choosy, well-mixed, and well-adapted generalists. And there are stretches in which the population evolves strong preference for one host species or the other, and performs well on that host species, and the other host species is nearly or completely unused.

**Figure 4.**
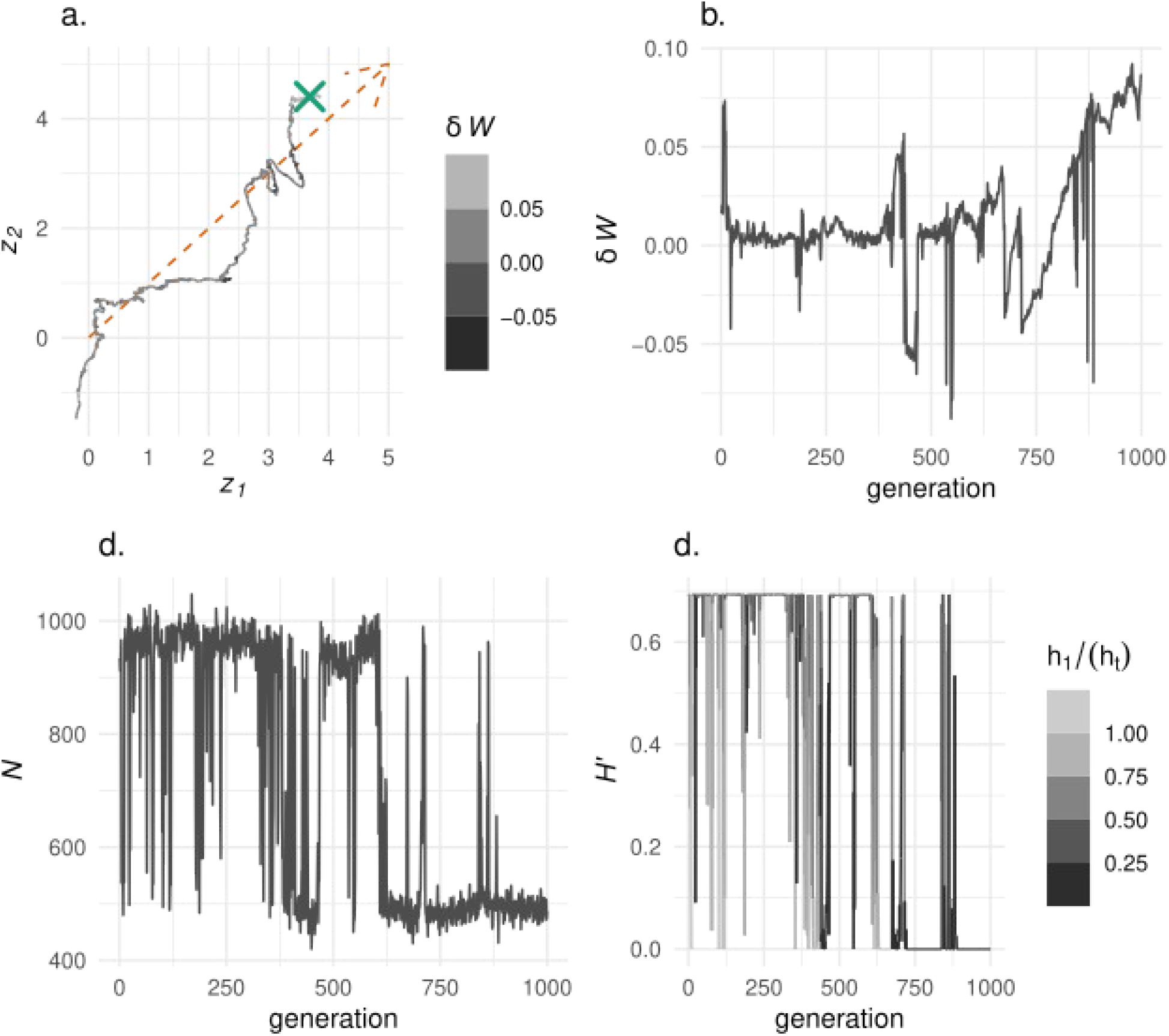
Example run under Model 2. Values for model parameters are *χ* = 1e-4, *ρ* = 1, *κ* = −0.9*σ^2^*/2, *M* = 2^−3^, and *δ_O_* = 0.005. (a) Changes in population mean values for performance traits (*z_1_* and *z_2_*), over 1000 generations after the burn-in period. The color of the tracing line changes with a population’s mean host-performance specificity; lighter shades of gray denote higher specificity. The dashed orange arrow shows the concomitant change in host performance optima (*O_j_*). The green ‘X’ denotes the population mean at the end of the simulation. Note the pronounced deviations of the populaiton’s evolutionary path from the optimal vector. The other three panels show the evolution of *δW* (b), *N* (c), and *H’* (d) following the burn-in period. In (d) the color of the tracing line changes with the proportion of the population found on *h_1_*; this allows us to see that the population temporarily specializes on different hosts.

We summarize our analysis of Model 2 with the plots in fig. 5. Here *χ* = 1e-4, and *ρ* = 1. In 5a-d, *μ* = 1e-7 and *δ_O_* = 0, that is, the optimal performance phenotype is the same in both host environments and does not change over the simulation. Without pleiotropy (*M* = 0) – in which case all preference mutations are conditionally neutral – simulated populations maintain high-performance generalism. With increasing preference-and-performance pleiotropy, populations spend more time as specialists on one host species or the other; *H’* declines steeply with increasing values for *M*, regardless of *κ*. (*δW* stays low, but given that performance optima are identical for each host species, this is as expected). Now look at figs. 5e-h, where *δ_O_* = 0.005 and *μ*=1e-9, that is, when the host-performance optima diverge, and we drop the rates of all kinds of mutation by two orders of magnitude. In this case, populations tend to have generalized host-use regardless of pleiotropy; even when *κ* = - 0.9*σ^2^/2* and *M* >> 1, *H’* drops only by about a third of its maximum value, and the mean value for *δW* across replications is < 0.2. So, the first set of simulations shows that with high absolute rates of mutation affecting preference and performance, populations tend to be host specialists, even when that rate is much lower than the rate of conditionally neutral mutation affecting performance, and when there are no differences in optimal performance phenotypes across hosts. Conversely, the second set of simulations show that when host environments do diverge, even when pleiotropy is more common than conditional neutrality, specialization can be prevented by reducing the absolute rate of pleiotropic mutation. With preference-and-performance pleiotropies, the preference component is key, and mutational pressure can drive arbitrary specialization across a homogeneous performance landscape.

**Figure 5.**
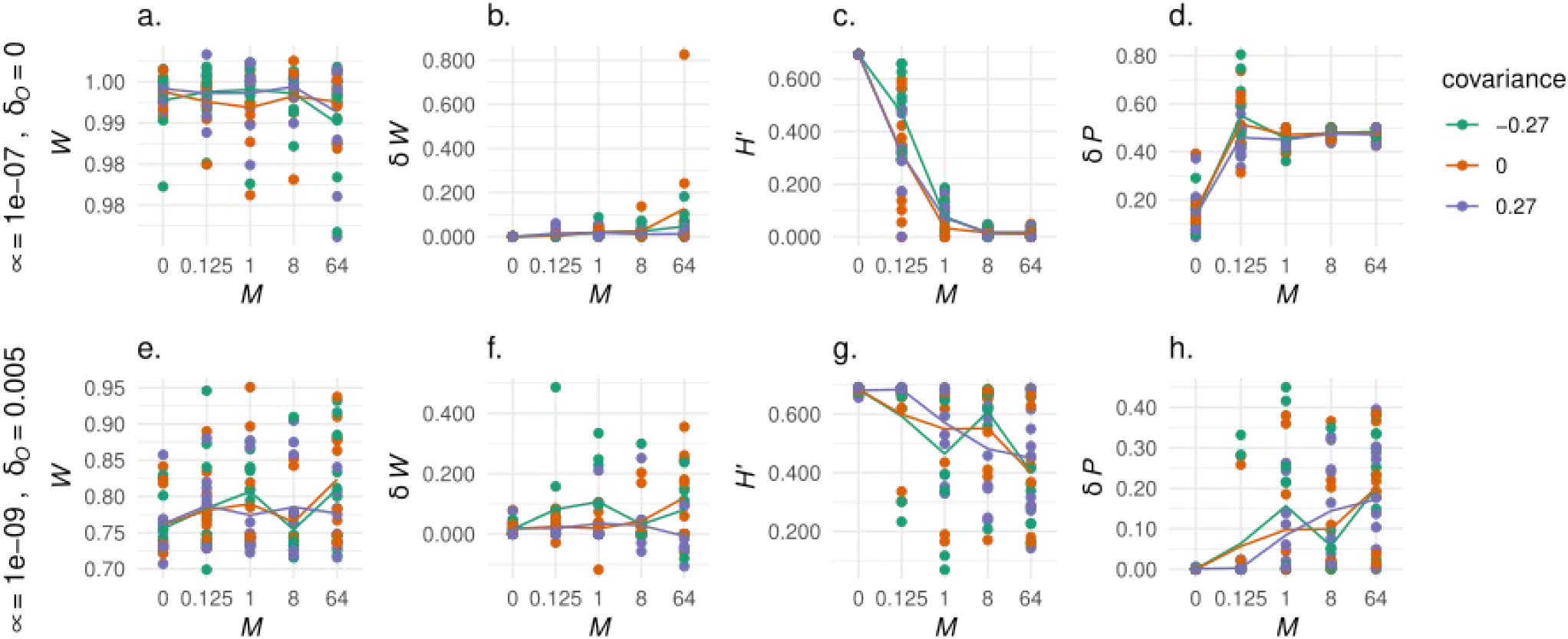
Summary of runs under Model 2. *M* is the ratio of pleiotropic to conditionally neutral mutation. Green lines are for pleiotropic covariances of −0.54, orange lines are for covariances of zero, and purple lines are for covariances of 0.54. All plots show mean population states over 1000 generations after the burn-in period. The top row (a–d) shows population states when *χ* = 1e-4, *ρ* = 1, *μ* = 1e-7 and *δ_O_* = 0, that is, host performance optima are fixed at the same value for both host species. The bottom row (e–h) is for simulations in which *χ* = 1e-4, *ρ* = 1, *μ* = 1e-9 *δ_O_* = 0.005, that is with slowly changing host performance optima, but reduced rates of mutation.

How then do we explain the pronounced evolutionary fluctuations of host-use breadth (e.g. fig. 4)? This is not something we observed in simulations under Model 1, but the occurrence of something like these fluctuations in nature has long been suspected (Janz and Nylin 2008; Hardy 2017). Here is how we see it. Negative density-dependent viability selection pushes for broad host-use. But the pressure of mutations affecting preference-and-performance pleiotropy can undermine such generalism. Moreover, local adaptation within a host environment is more efficient when the population evolves a strong host preference (Whitlock 1996), and thus, a substantial lag of mean populations phenotypes behind host-performance optima can promote specialization. To test this intuition, we performed timeseries regression analyses of 20 individual simulations with *χ*=1e-4, *ρ*=0.1, *δ_O_*=0.005, *M*=2^−3^, *κ* = - 0.9*σ^2^/*2, *μ*=1e-7. For our response variable we define an index *L* of the rate of adaptation that is a function of the extent to which mean performance values lagged behind host optima. Taking advantage of the fact that for each simulation there is a bimodal distribution for *H’*, for each generation, we classify a population as being in either a specialist or generalist state, using a cut-off value for *H’* of 0.2 When a population is in the specialist state, we define *L* as 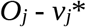, where 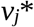 is the population mean. When the population is in the generalist state, *L* is 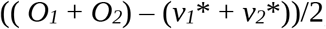, that is, the average lag between host optima and mean performance values. As predictor variables we used *N*, *H’*, and *H*. Using the R package *metafor* (Viechtbauer and Viechtbauer, 2015) to perform a simple meta-analysis, with the estimated effects of *H’* on *L* as the effect size, and weighting effect sizes by standard errors, we found that average effect is positive and significant (effect = 0.52, p-value < 0.001); as expected, host-use generalists are slower to adapt to diverging host-use optima.

As optimal performance phenotype values in each host environment move farther away from a population’s starting genotype values, it takes more alleles to produce a locally optimal phenotype (Holt et al. 2003), and the potential downside grows for recombination in the offspring of parents born in different host environments. Likewise as the proportion of pleiotropic mutation increases, it becomes harder to optimize performance in both host environments without evolving strong preference for one or the other. Thus, large displacements of *O_1_* and *O_2_* from a population’s initial mean performance genotype values, along with more pervasive pleiotropy increase the time a population spends as a specialist.

Are preference-and-performance pleiotropies likely? One possibility would be a mutation at a site that regulates the expression of preference and performance genes. Another would be a mutation at a site encoding an enzyme involved in host-sensing, affecting feeding rates during development and oviposition as an adult. Yet another would be a gene for an enzyme involved in both host-sensing (e.g., degrading a host secondary compound within a sensillum on an instect’s hypopharynx), and detoxification (degrades the same compound in the insect’s gut). In any case, the importance of pleiotropy for the evolution of niche specialization may depend on the natural occurrence of mutations that directly affect both habitat-dependent performance and assortative mating. In that respect, our inferences resonate with the sympatric speciation research; specialization is much more likely if there are “magic alleles” that promote local adaptation in a way that directly impedes gene flow across demes in a structured metapopulation (Bolnick and Fitzpatrick, 2007). But note that in our models the “magic alleles” that affect host-use preference as well as performance do not promote species divergence *per se*; they cause niche contraction rather than segmentation.

A few words about quantitative pleiotropy might help to contextualize our study. In classical single-locus population genetic models, alleles have direct fitness effects, and by definition an antagonistically pleiotropic allele improves fitness – or preference – for one environment and decreases it in another (Templeton 2021; e.g., Levene 1953; Rausher 1984). In contrast, in a quantitative genetics framework, alleles indirectly affect fitness via their effects on quantitative traits and the matching of trait values to environmental optima (e.g., Burger & Lynch 1995; Holt et al. 2002). Furthermore, an allele’s pleiotropic effects can be positive or antagonistic depending on the genetic environment, that is, the sum of all other allele effects across QTLs. In a word, polygenic pleiotropy is relative. Therefore, we cannot use quantitative genetics to test the predictions of the single-locus theory; that theory hinges on a concept of pleiotropy that has no direct parallel in quantitative genetics. On the other hand, since herbivorous insect host-use phenotypes are polygenic and quantitative, quantitative genetics can give us a better sense for how pleiotropy actually works in nature.

## Conclusions

It would seem that pleiotropy can indeed drive quantitative niche specialization, especially when pleiotropic alleles affect preference as well as performance. On the other hand, if pleiotropies affect only performance, even with freely-evolvable habitat preference, to drive specialization they need to be much more common and antagonistic than has been observed in nature. And if habitats change quickly, or differ enough in productivity, quantitative specialization can readily evolve without pleiotropy. So, pleoiptropy is just one among the many factors that can lead to habitat specialization.

We are not the first to theoretically demonstrate that specialization can evolve without simple genetic trade-offs (e.g., Kawecki 1994; Fry 1996; Whitlock 1996; Kawecki et al. 1997; Dragi 2021). But assuming monogeny, the joint evolution of niche preference and performance has seemed an especially powerful mechanism (Ravigne et al. 2007). Our models show under more realistic polygenic architectures, the influence of pleiotropy is more limited. And so we suggest that future theoretical and experimental work might look deeper into that extent of pleiotropy on habitat preference and performance, and more broadly for what stands in the way of the evolution of generalism (Hardy et al., 2020).

---

**Fig S1. Pleiotropic allele effects on performance.** In Model 1, pleiotropic allele effects on host-use performance are drawn from a bivariate normal distribution. Here we show 100 random deviates from a distribution with means of zero, variances of 0.6, and covariences of 0.54. This is one of the three distributions used in our simulations. General vigor alleles are shown with a “+”, General weakness alleles with a “o”, and antagonistically plieotropic alleles with an “x.” In this case, most of the sampled alleles are antagonistically pleiotropic. At the start of each simulation, the optimal performance phenotype value in each host environment is zero. Then, after the burn-in period *O_j_* increases by *δ_O_* each generation.

**Figure S2. Summary of runs under Model 1, with *ρ*=0.1.** Green lines are for pleiotropic covariances of −0.54, orange lines are for covariances of zero, and purple lines are for covariances of 0.54. All plots show mean population states over 1000 generations after the burn-in period. The left column (a–d) shows population states when host species are equally productive (*K_1_=K_2_*), and in which optimal performance values change slowly (*δ_O_*=0.005). The middle column (e–h) is for simulations in which *K_1_=K_2_*, and optimal performance value change rapidly (*δ_O_*=0.1). The right column (i–l) is for simulations in which *K_1_<K_2_ (pK_1_* = 0.25) and optimal performance values change slowly.

**Figure S3. Summary of runs under Model 1, with *χ*=1e-6.** Green lines are for pleiotropic covariances of −0.54, orange lines are for covariances of zero, and purple lines are for covariances of 0.54. All plots show mean population states over 1000 generations after the burn-in period. The left column (a–d) shows population states when host species are equally productive (*K_1_*=*K_2_*), and in which optimal performance values change slowly (*δ_O_*=0.005). The middle column (e–h) is for simulations in which *K_1_=K_2_*, and optimal performance value change rapidly (*δ_O_*=0.1). The right column (i–l) is for simulations in which *K_1_<K_2_ (pK_1_* = 0.25) and optimal performance values change slowly.

## Notes

https://github.com/n8-rd/PleioPoly

### Competing Interest Statement

The authors have declared no competing interest.

### Summary of Updates

This version is a major update from the last. We have changed several core model components, such as the implementation of pleiotropy and the evolution of host preference. We explore a broader range of parameter values, for example, the strength of selection and the rate of recombination. And we place our work in the context of previous work on the evolution of niche specificity and sympatric speciation.

