## Supplementary figures and images for "Niche specificity, polygeny, and pleiotropy in herbivorous insects"

### Supplementary figure 1

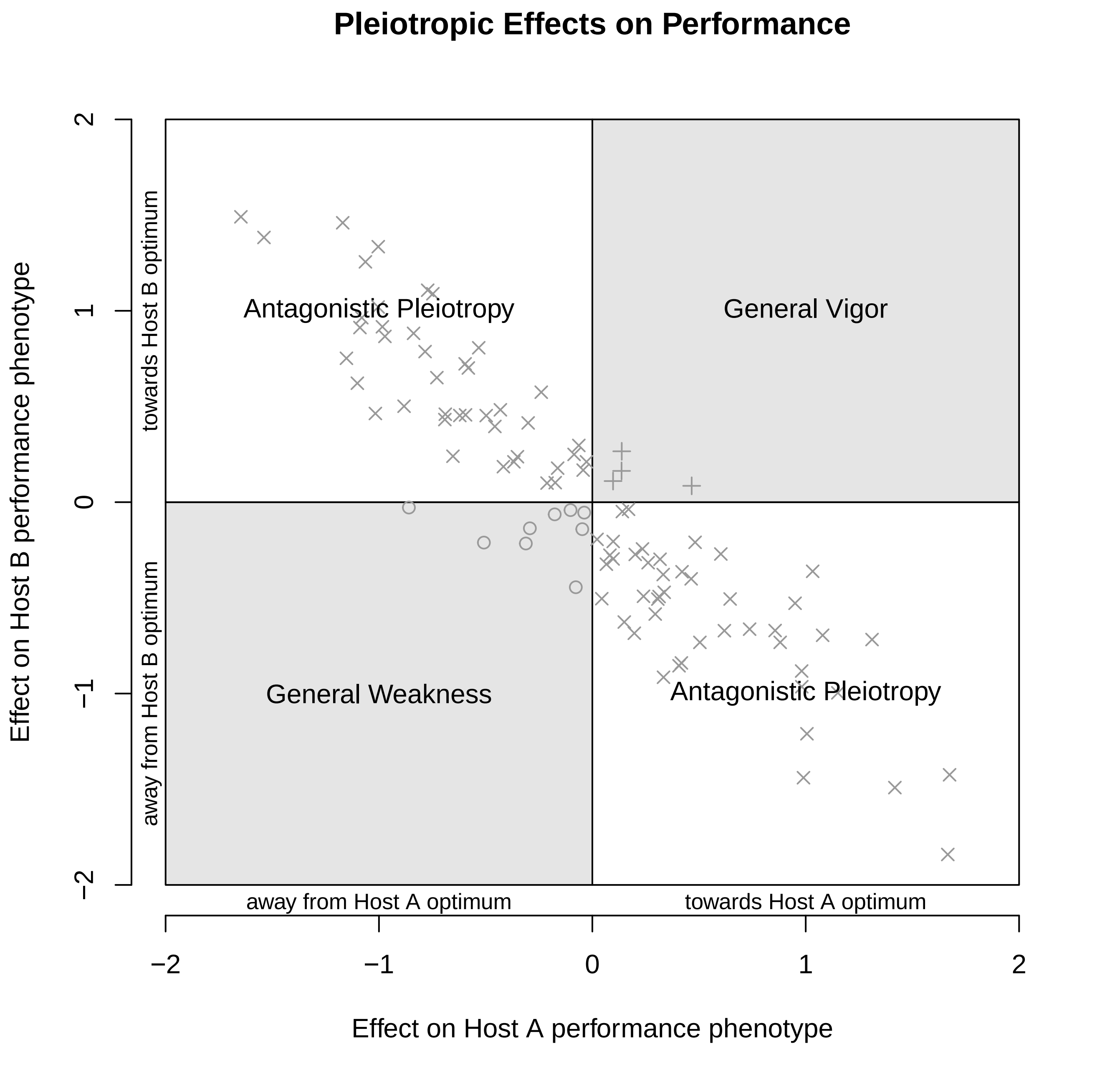

### Supplementary figure 2

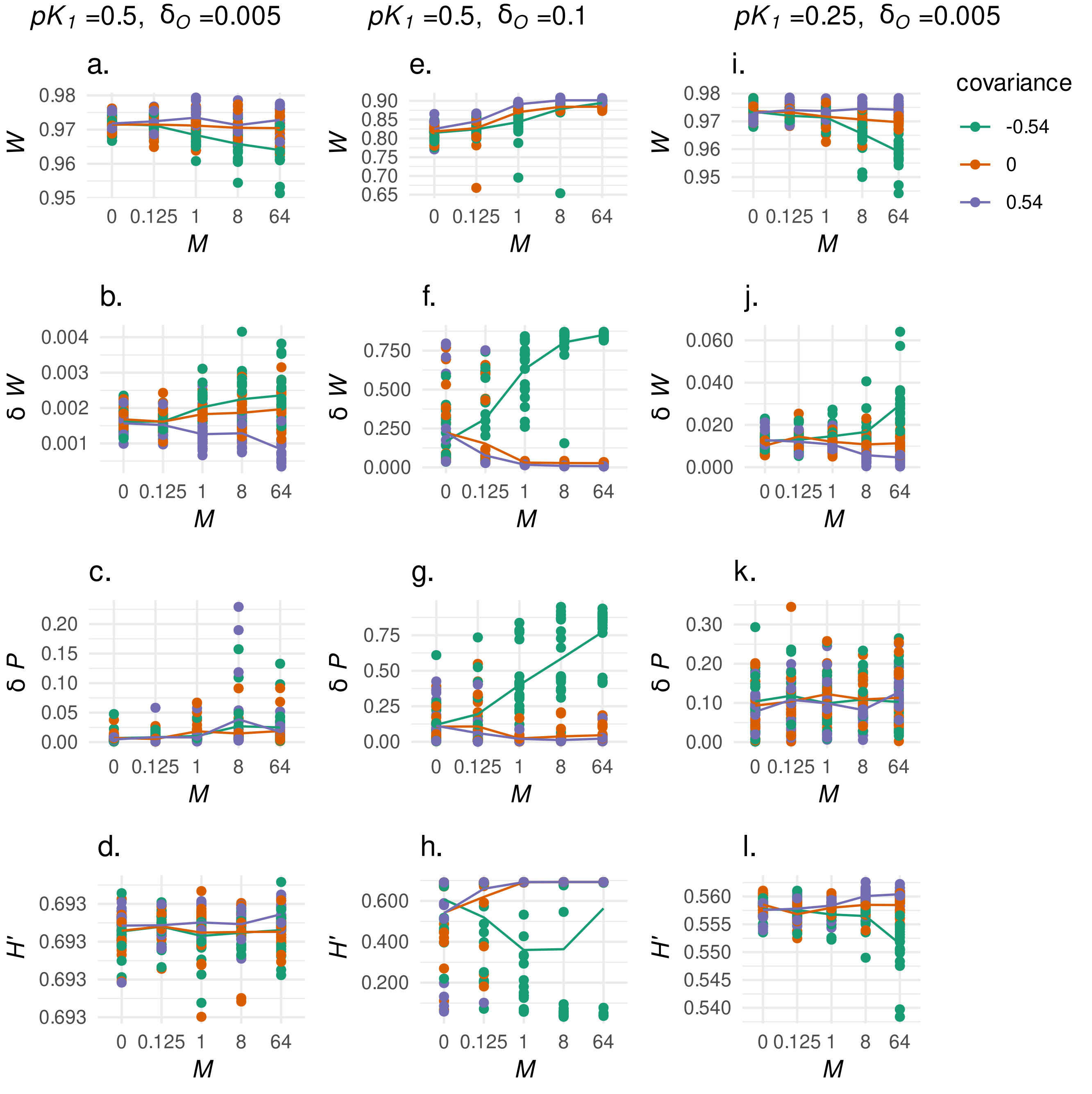

### Supplementary figure 3

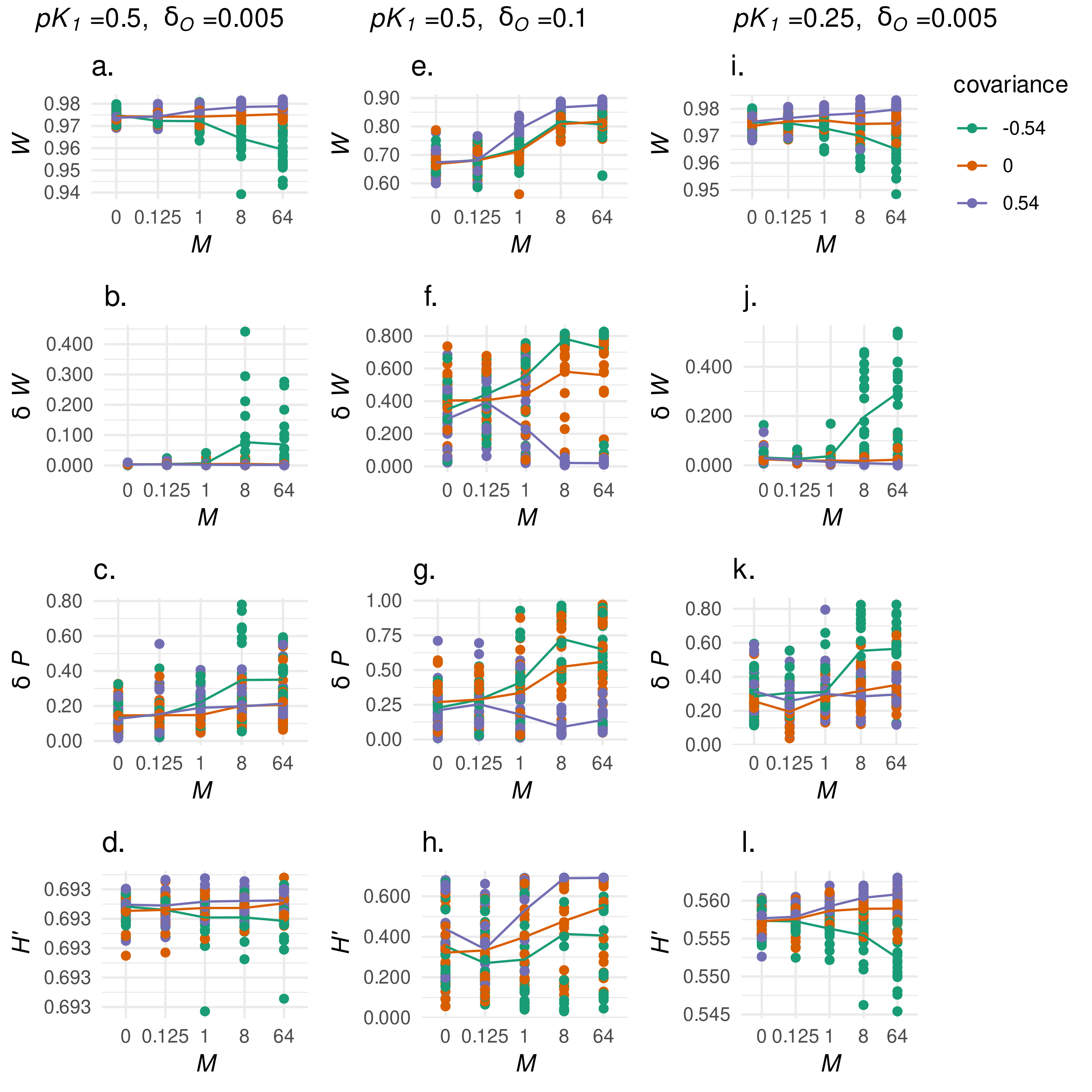
